## Supplemental Information for "Widespread contribution of transposable elements to the rewiring of mammalian 3D genomes and gene regulation"

### Supplementary Information

#### Supplementary Figure 1

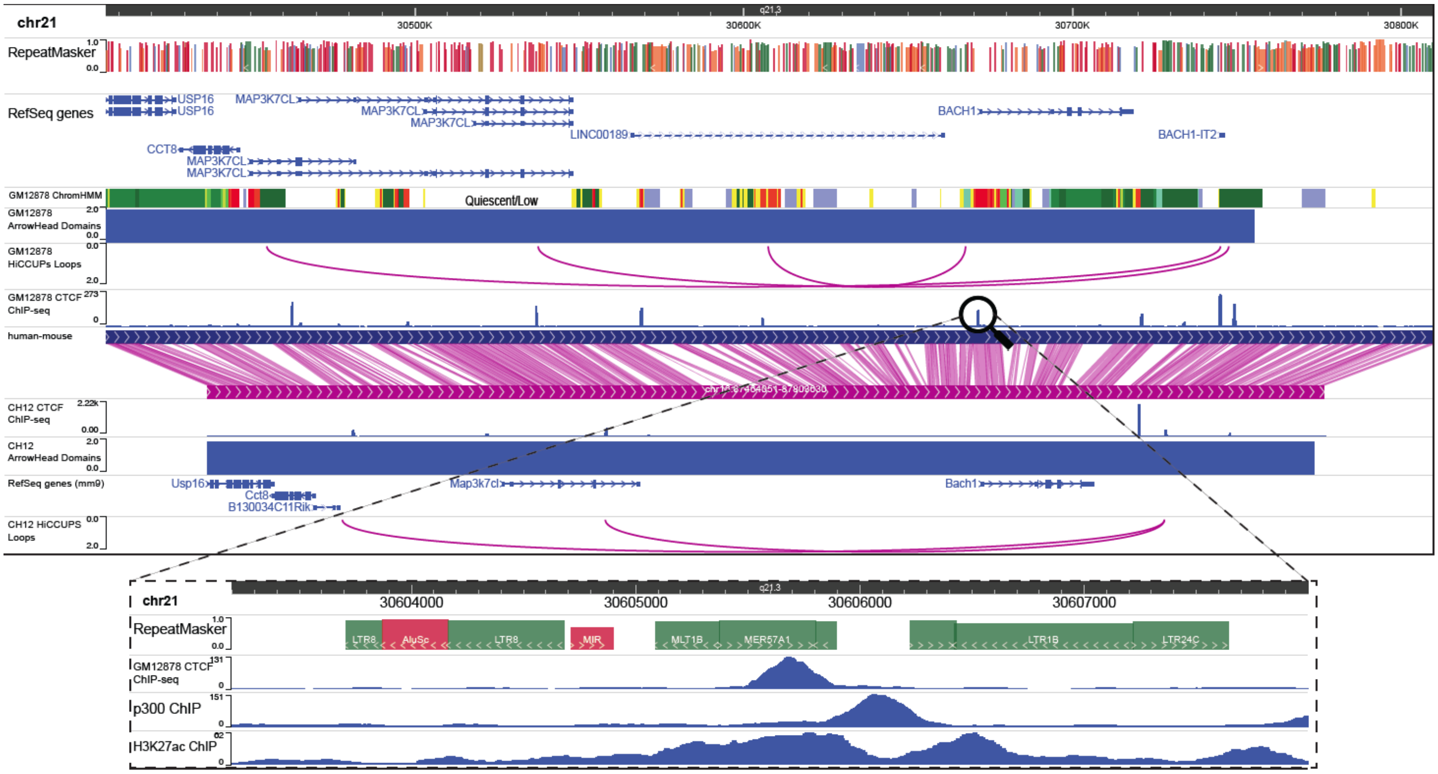


Genome browser screenshot displaying the MER57A1 candidate and the local histone, binding, 3D landscape in human (main top) and mouse (main bottom). Zoomed in view of the MER571A1 candidate with CTCF, p300, and H3K27ac ChIP-seq peaks (inset).

#### Supplementary Figure 2

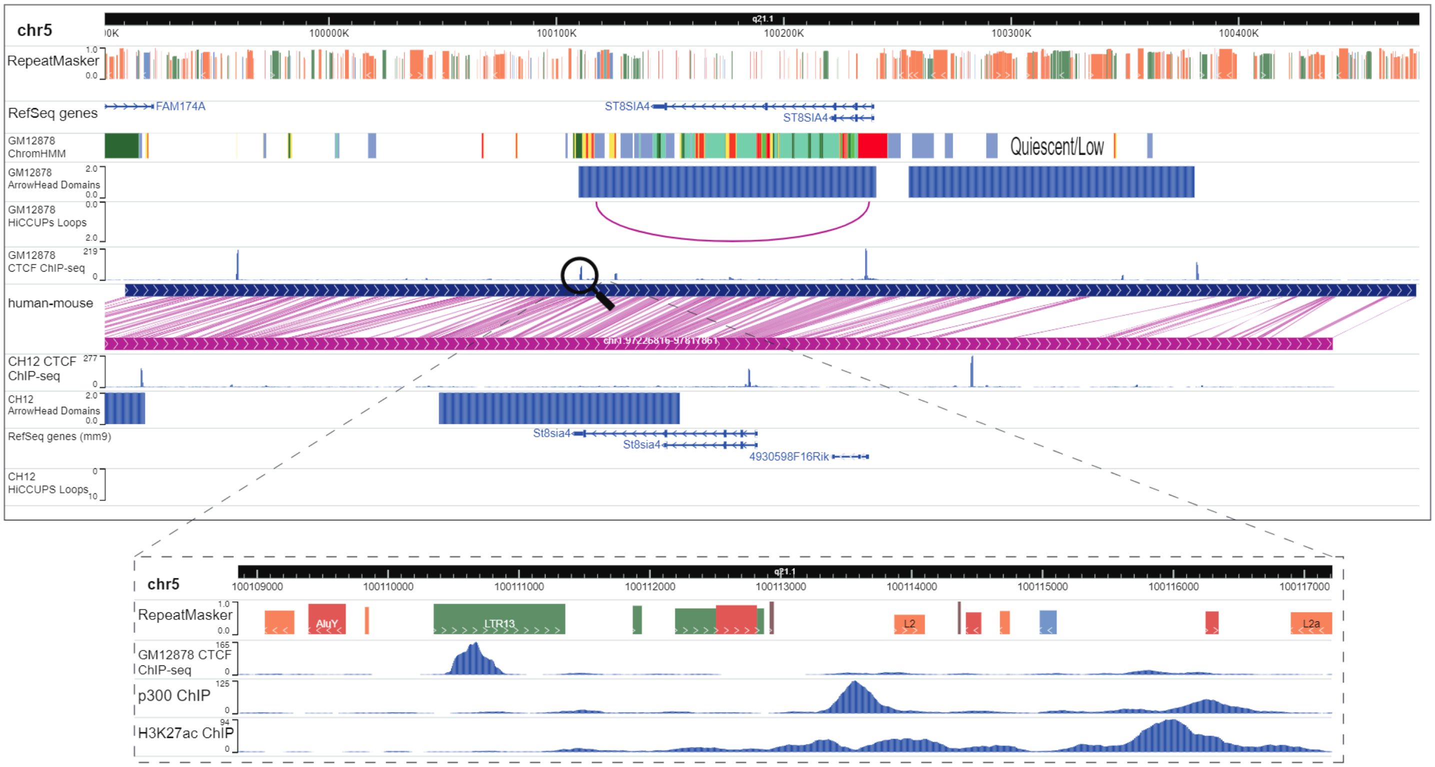


Genome browser screenshot displaying the LTR13 candidate and the local histone, binding, 3D landscape in human (main top) and mouse (main bottom). Zoomed in view of the LTR13 candidate with CTCF, p300, and H3K27ac ChIP-seq peaks (inset).

**Supplementary Table 1: L1MC1-KO1 HiC² Library Statistics**

| | |
|---|---|
| Sequenced Read Pairs | 101,630,014 |
| Normal Paired | 90,448,638 (89.00%) |
| Chimeric Paired | 4,407,936 (4.34%) |
| Chimeric Ambiguous | 3,513,938 (3.46%) |
| Unmapped | 3,259,502 (3.21%) |
| Ligation Motif Present | 29,354,231 (28.88%) |
| Alignable (Normal+Chimeric Paired) | 94,856,574 (93.34%) |
| Unique Reads | 28,973,498 (28.51%) |
| PCR Duplicates | 65,831,160 (64.78%) |
| Optical Duplicates | 51,916 (0.05%) |
| Library Complexity Estimate | 30,299,572 |
| Intra-fragment Reads | 325,061 (0.32% / 1.12%) |
| Below MAPQ Threshold | 19,264,376 (18.96% / 66.49%) |
| Hi-C Contacts | 9,384,061 (9.23% / 32.39%) |
| Ligation Motif Present | 2,054,781 (2.02% / 7.09%) |
| 3' Bias (Long Range) | 69% - 31% |
| Pair Type %(L-I-O-R) | 25% - 25% - 25% - 25% |
| Inter-chromosomal | 2,431,679 (2.39% / 8.39%) |
| Intra-chromosomal | 6,952,382 (6.84% / 24.00%) |
| Short Range (<20Kb) | 2,615,029 (2.57% / 9.03%) |
| Long Range (>20Kb) | 4,337,335 (4.27% / 14.97%) |

**Supplementary Table 2: L1MC1-KO47 HiC² Library Statistics**

| | |
|---|---|
| Sequenced Read Pairs | 29,681,164 |
| Normal Paired | 26,419,325 (89.01%) |
| Chimeric Paired | 1,505,030 (5.07%) |
| Chimeric Ambiguous | 835,886 (2.82%) |
| Unmapped | 920,923 (3.10%) |
| Ligation Motif Present | 8,772,298 (29.56%) |
| Alignable (Normal+Chimeric Paired) | 27,924,355 (94.08%) |
| Unique Reads | 15,224,433 (51.29%) |
| PCR Duplicates | 12,685,207 (42.74%) |
| Optical Duplicates | 14,715 (0.05%) |
| Library Complexity Estimate | 20,445,316 |
| Intra-fragment Reads | 166,830 (0.56% / 1.10%) |
| Below MAPQ Threshold | 6,958,217 (23.44% / 45.70%) |
| Hi-C Contacts | 8,099,386 (27.29% / 53.20%) |
| Ligation Motif Present | 2,199,042 (7.41% / 14.44%) |
| 3' Bias (Long Range) | 71% - 29% |
| Pair Type %(L-I-O-R) | 25% - 25% - 25% - 25% |
| Inter-chromosomal | 1,929,082 (6.50% / 12.67%) |
| Intra-chromosomal | 6,170,304 (20.79% / 40.53%) |
| Short Range (<20Kb) | 2,227,463 (7.50% / 14.63%) |
| Long Range (>20Kb) | 3,942,828 (13.28% / 25.90%) |

**Supplementary Figure 3: Virtual 4C Analysis of Downstream Enhancers**

#### Virtual 4C Analysis of Downstream Enhancers

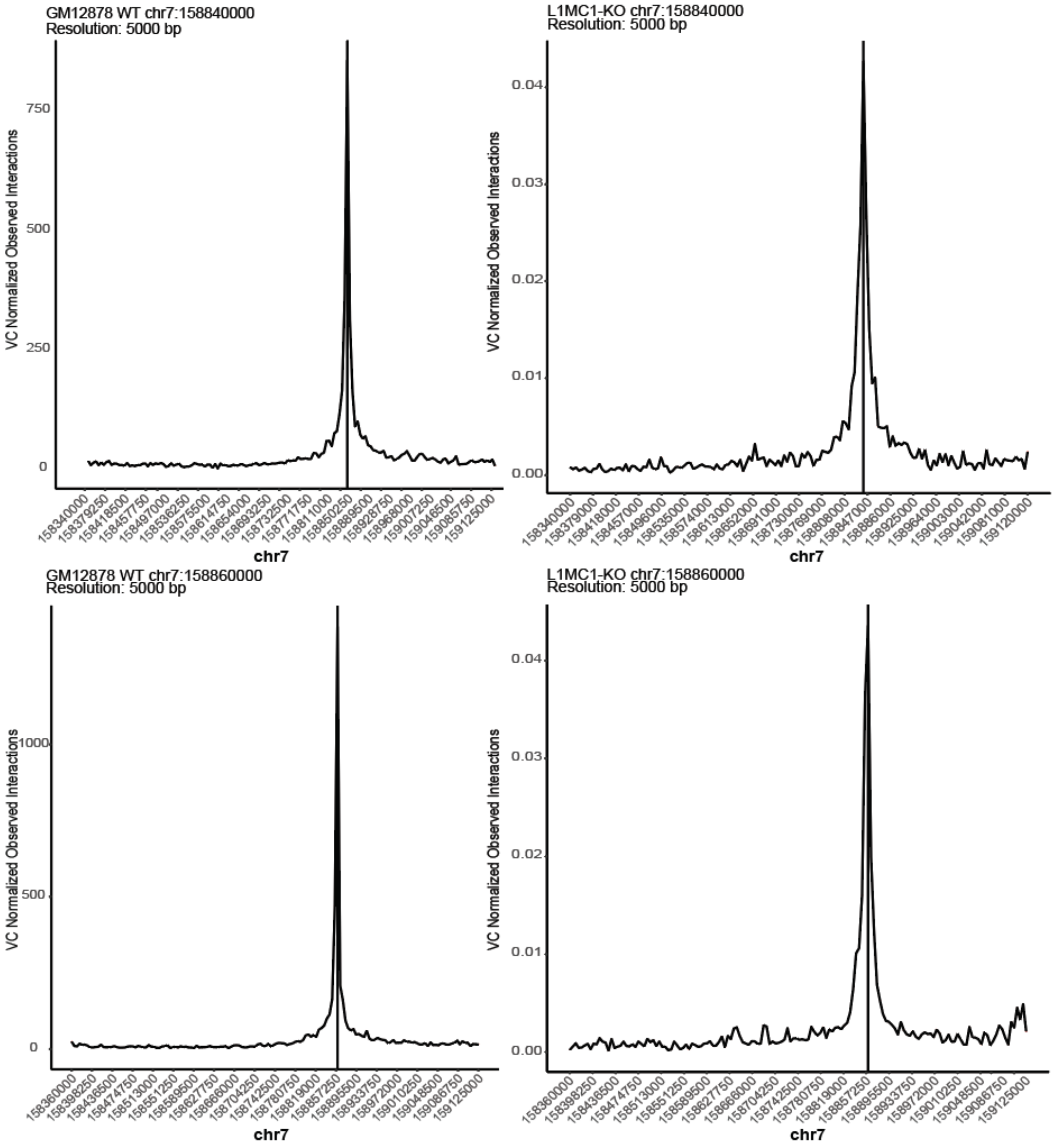


Virtual 4C analysis showing interactions between downstream enhancers and active domain in wildtype and L1MC1-KO lines.

**Supplementary Table 3: Loop and TAD Call Data Sources**

Data sources of loop and TAD calls used in all analyses.

| Species | Structure | Assay | Cell Type | Link |
|---|---|---|---|---|
| Human | TADs | Hi-C | HSC_aml | https://www.biorxiv.org/content/10.1101/2020.04.18.047738v1 |
| Human | TADs | Hi-C | HSC_aml | |
| Human | TADs | Hi-C | HSC_aml | |
| Human | TADs | Hi-C | HSC | |
| Human | TADs | Hi-C | HSC | |
| Human | TADs | Hi-C | HSC | |
| Human | Loops | Hi-C | HSC_aml | |
| Human | Loops | Hi-C | HSC_aml | |
| Human | Loops | Hi-C | HSC_aml | |
| Human | Loops | Hi-C | HSC | |
| Human | Loops | Hi-C | HSC | |
| Human | Loops | Hi-C | HSC | |
| Human | Loops | H3K4me2_ChIA-PET | CD4+ T Cells (CD4) | https://www.nature.com/articles/cr201215 |
| Human | TADs | Hi-C | Cortex | https://www.nature.com/articles/nature11082 |
| Human | TADs | Hi-C | hESC | |
| Human | TADs | Hi-C | IMR90 | |
| Human | Loops | H3K27ac_HiChIP | GM12878 | https://www.nature.com/articles/ng.3963 |
| Human | Loops | H3K27ac_HiChIP | HCASMC | |
| Human | Loops | H3K27ac_HiChIP | K562 | |
| Human | Loops | H3K27ac_HiChIP | MyLa | |
| Human | Loops | H3K27ac_HiChIP | Naive_T | |
| Human | Loops | H3K27ac_HiChIP | TH17 | |
| Human | Loops | H3K27ac_HiChIP | TReg | |
| Mouse | Loops | H3K27ac_HiChIP | MES | |
| Human | Loops | CTCF_HiChIP | GM12878 | https://www.nature.com/articles/nmeth.3999#Sec19 |
| Mouse | Loops | CTCF_HiChIP | mESC | |
| Human | TADs | Hi-C | hESC | https://www.nature.com/articles/s41586-019-1812-0 |
| Human | TADs | Hi-C | hESC | |

| | | | | |
|---|---|---|---|---|
| Human | TADs | Hi-C | hESC | |
| Human | TADs | Hi-C | hESC | |
| Human | TADs | Hi-C | hESC | |
| Mouse | TADs | Hi-C | mESC | |
| Mouse | TADs | Hi-C | mESC | |
| Mouse | TADs | Hi-C | mESC | |
| Human | TADs | Hi-C | Lymphoma | https://www.nature.com/articles/s41588-018-0338-y |
| Human | TADs | Hi-C | Lymphoma | |
| Dog | TADs | Hi-C | Liver | https://www.ncbi.nlm.nih.gov/pmc/articles/PMC4542312/ |
| Human | TADs | Hi-C | RPE1 | https://www.ncbi.nlm.nih.gov/pmc/articles/PMC4978254/#d36e641 |
| Human | TADs | Hi-C | RPE1 | |
| Human | TADs | Hi-C | RPE1 | |
| Mouse | TADs | Hi-C | PATSKI | |
| Mouse | TADs | Hi-C | mESC | |
| Rhesus Macaque | TADs | Hi-C | Fibroblasts | |
| Human | Loops | Hi-C | RPE1 | |
| Human | Loops | Hi-C | RPE1 | |
| Human | Loops | Hi-C | RPE1 | |
| Mouse | Loops | Hi-C | PATSKI | |
| Rhesus Macaque | Loops | Hi-C | Fibroblasts | |
| Mouse | TADs | Hi-C | G1E-ER4 | https://www.ncbi.nlm.nih.gov/pmc/articles/PMC5393350/ |
| Mouse | TADs | Hi-C | G1E-ER4 | |
| Human | TADs | Hi-C | Adrenal Cells (AD) | https://www.ncbi.nlm.nih.gov/pmc/articles/PMC5478386/ |
| Human | TADs | Hi-C | Aorta (AO) | |
| Human | TADs | Hi-C | Bladder (BL) | |
| Human | TADs | Hi-C | Cortex (CO) | |
| Human | TADs | Hi-C | GM12878 | |
| Human | TADs | Hi-C | hESC (H1) | |
| Human | TADs | Hi-C | Hippocampus (HC) | |
| Human | TADs | Hi-C | IMR90 | |
| Human | TADs | Hi-C | Lung (LG) | |
| Human | TADs | Hi-C | Liver(LI) | |

| | | | | |
|---|---|---|---|---|
| Human | TADs | Hi-C | Left Ventrical (LV) | |
| Human | TADs | Hi-C | Mesendoderm (MES) | |
| Human | TADs | Hi-C | Mesenchymal (MSC) | |
| Human | TADs | Hi-C | NPC | |
| Human | TADs | Hi-C | Ovary (OV) | |
| Human | TADs | Hi-C | Pancreas (PA) | |
| Human | TADs | Hi-C | Psoas Muscle (PO) | |
| Human | TADs | Hi-C | Right Ventrical (RV) | |
| Human | TADs | Hi-C | Small Bowel (SB) | |
| Human | TADs | Hi-C | Spleen (SX) | |
| Human | TADs | Hi-C | Trophoblast-like (TRO) | |
| Human | Loops | Hi-C | THP-1_pma | https://www.ncbi.nlm.nih.gov/pmc/articles/PMC5610110/ |
| Human | Loops | Hi-C | THP-1 | |
| Mouse | TADs | Hi-C | AML12 | https://www.ncbi.nlm.nih.gov/pmc/articles/PMC5793783/ |
| Mouse | Loops | Hi-C | mESC | https://www.ncbi.nlm.nih.gov/pmc/articles/PMC6327227/ |
| Mouse | Loops | Hi-C | NSC | |
| Human | TADs | Hi-C | LCL | https://www.ncbi.nlm.nih.gov/pmc/articles/PMC6883528/ |
| Human | Loops | CTCF_ChIA-PET | Consensus | |
| Mouse | TADs | Hi-C | CH12-LX | https://www.ncbi.nlm.nih.gov/pubmed/25497547 |
| Human | TADs | Hi-C | GM12878 | |
| Human | TADs | Hi-C | HeLa | |
| Human | TADs | Hi-C | HMEC | |
| Human | TADs | Hi-C | HUVEC | |
| Human | TADs | Hi-C | IMR90 | |
| Human | TADs | Hi-C | K562 | |
| Human | TADs | Hi-C | KBM7 | |
| Human | TADs | Hi-C | NHEK | |
| Human | Loops | Hi-C | GM12878 | |
| Human | Loops | Hi-C | HeLa | |
| Human | Loops | Hi-C | HMEC | |

| | | | | |
|---|---|---|---|---|
| Human | Loops | Hi-C | HUVEC | |
| Human | Loops | Hi-C | IMR90 | |
| Human | Loops | Hi-C | K562 | |
| Human | Loops | Hi-C | KBM7 | |
| Human | Loops | Hi-C | NHEK | |
| Mouse | Loops | Hi-C | CH12-LX | |
| Human | TADs | Hi-C | HAP1 | https://www.pnas.org/content/112/47/E6456.long |
| Human | Loops | Hi-C | HAP1 | |
| Human | Loops | Hi-C | HSC | https://www.sciencedirect.com/science/article/pii/S1097276520302604 |
